# On the relationship between epistasis and genetic variance-heterogeneity

**DOI:** 10.1101/119727

**Authors:** Simon Forsberg, Örjan Carlborg

## Abstract

Epistasis and genetic variance heterogeneity are two non-additive genetic inheritance patterns that are often, but not always, related. Here we use theoretical examples and empirical results from analyses of experimental data to illustrate the connection between the two. This includes an introduction to the relationship between epistatic gene-action, statistical epistasis and genetic variance heterogeneity and a brief discussion about how other genetic processes than epistasis can also give rise to genetic variance heterogeneity.

**Highlight:** Genetic effects on the trait variance, rather than the mean, have been found in several studies. Here we discuss how this sometimes, but not always, can be caused by epistasis.

## Introduction

Complex traits are generally affected by many alleles at different loci throughout the genome. A central question when trying to understand the genetic mechanisms regulating a complex trait is therefore: do the different alleles act independently of each other, or are the phenotypic effects of certain alleles dependent on the genetic background at other loci? Such dependencies between alleles at different loci are referred to as genetic interactions, or epistasis.

In quantitative genetics, the phenotypic variance (V_P_) is typically partitioned into one component (V_G_) that is due to genetic, and another component (V_E_) that is due to non-genetic, influences on the trait variance. The genetic variance (V_G_) can then be further partitioned into additive (V_A_), dominance (V_D_), and epistatic (V_E_) variance components^1^. This variance partitioning forms the basis of the metric heritability, which is defined as 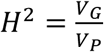(broad sense heritability), or 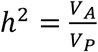(narrow sense heritability). The concept of additive genetic variance has been very useful in animal and plant breeding efforts^1^, since V_A_ captures the “breedable” genetic contributions to the resemblance between relatives. The epistatic variance, however, is of little value in breeding and was originally introduced as a nuisance parameter in the genetic model^1^. These genetic variance components are population level metrics, specific to a certain population, and they do not have a straightforward functional interpretation in terms of gene action^2,3^. Epistatic gene action, i.e. when the effect of an allele at one locus varies depending on the genotype at another locus, is therefore not directly proportional to the level of epistatic variance in a population. This as it will usually contribute to both the additive and epistatic genetic variances^4–6^. To which extent epistatic gene action will contribute additive genetic variance is determined by allele frequencies, the type of genetic interactions, and how the genetic models used are parameterized^3,5,7^. To clarify the distinction between gene action and genetic variance, it has sometimes been highlighted that additive genetic variance can be an “emergent property” of non-additive gene action. We refer readers interested in this topic to previous work^2–7^, as our focus here is on the connection between epistatic gene action and genetic variance-heterogeneity.

### Genetic regulation of the variability of a trait

The concept that also the trait variability could be under direct genetic control was introduced already in the 1940s when Waddington presented the idea of canalization, where he suggested that natural selection could act to produce traits that are robust to environmental and genetic perturbations^8^. He partly based his ideas on the observation that natural populations often are less variable than artificial populations of the same species. More recently, *Hill and Mulder^9^* proposed that “the environmental variation” can be regarded as a phenotype in its own right. One can then invoke much of the quantitative genetics methodology to search for genetic determinants of this phenotype. They consequently called this phenomenon “genetic control of the environmental variation”, a terminology which implies that it is the randomness, or instability, of the trait that is genetically controlled. Several studies have recently mapped individual loci where the different alleles affect not only the mean, but also the variance of traits^9–11^. These loci can be detected since the variability of the measured trait differs between groups of individuals that carry alternative alleles at the locus. A simple example would be two groups of humans, where the group of individuals homozygote for a certain allele include both very short and very tall individuals, while the second group that is homozygote for the alternative allele include individuals of similar height. This would lead to genetic variance heterogeneity between the two groups of individuals. Note that the mean height does not have to be different between the groups in order for this to occur (Fig. 1).

**Figure 1.**
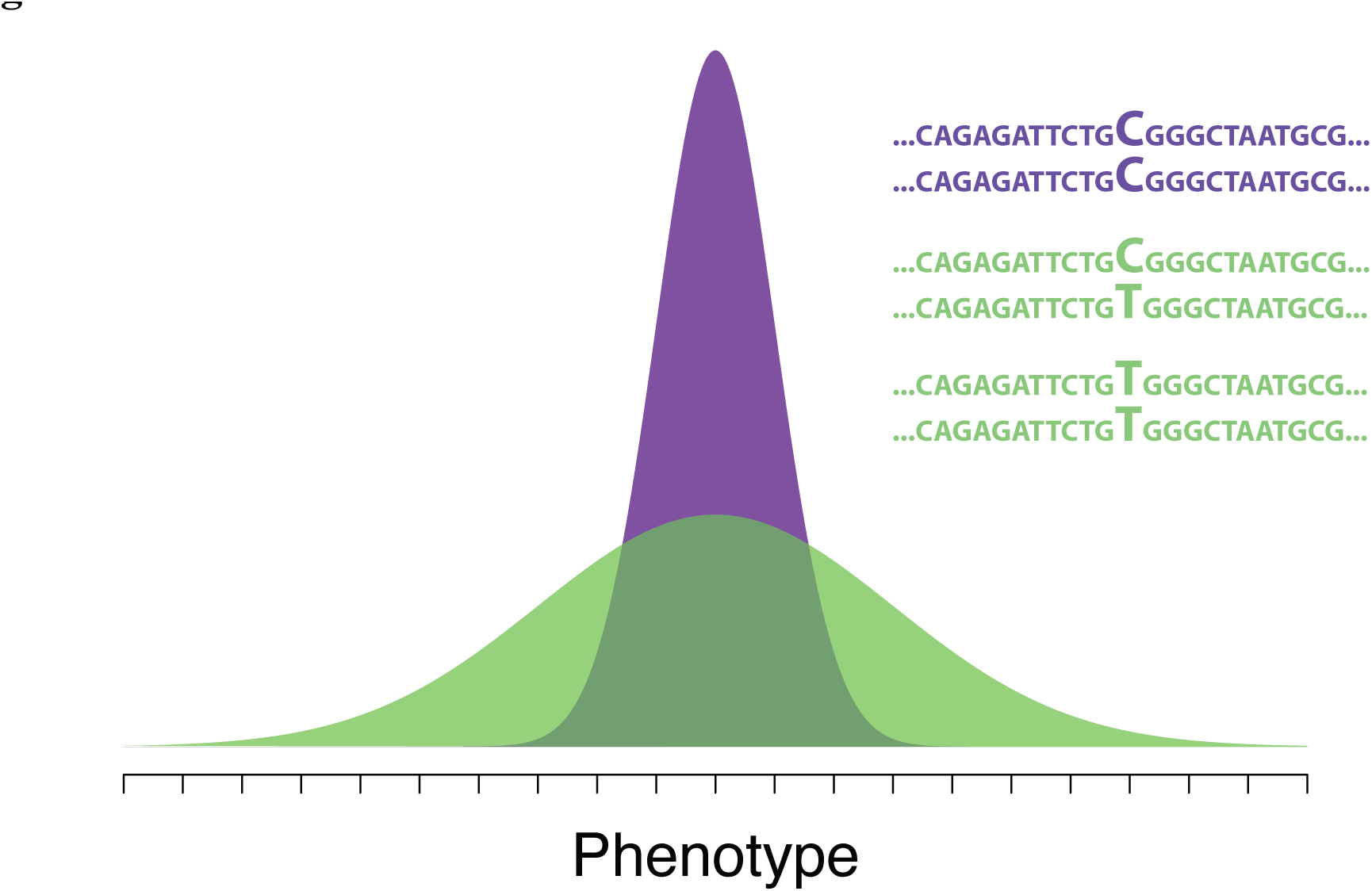
Illustration of the phenotypic distributions for the three genotypes at a locus displaying genetic variance heterogeneity. Individuals with the genotype C/C display a narrow phenotypic distribution, whereas individuals with the genotypes C/T and T/T display more variable phenotypes. There is no mean difference between the two genotypes, and although the genetic effect of the locus influences the total phenotypic variance in the population, it does not contribute any additive genetic variance and consequently does not contribute to the narrow-sense heritability for the trait.

A growing body of evidence thus suggests that phenotypic variability can be genetically controlled^10,12,13^. As genetic variance heterogeneity is a statistical measure defined on the population level it is, in the same way as additive and epistatic genetic variance, a property that could emerge from several different underlying genetic mechanisms. A number of different genetic mechanisms that in theory can lead to genetic variance heterogeneity have been proposed and discussed in the literature^11,13,14^. Consequently, it is not possible to make immediate functional interpretations of genetic variance heterogeneity associations. The underlying mechanisms revealed to date, however, have one thing in common: they reach beyond the assumption of bi-allelic loci affecting the trait mean, which is the mode of inheritance typically assumed in quantitative genetics studies. We will here first use two recent empirical dissections of variance-heterogeneity associations to illustrate the underlying mechanisms revealed in these studies: alleles increasing stochasticity of a trait and linkage-disequilibrium patterns across alleles at linked loci. After that, we will proceed to a third mechanism that can lead to genetic variance heterogeneity, and which is the main topic of this paper, genetic interactions or epistasis.

### Individual alleles can have a direct effect on the variability of a trait

Genetic variance heterogeneity is a somewhat abstract concept, since variance only has a meaning for groups, not for individuals. What does it mean that a certain allele increases the variance of a trait? One way to illustrate this is by using the results from *Ayroles et al*^12^, a study of variability in locomotor handedness in fruit flies. In this study, it was demonstrated that the degree of variability in how flies turn left and right in a Y-shaped maze was heritable. Further, some lines of flies displayed high levels of intragenotypic variability among individuals, whereas other lines had a low variability. This difference could in part be explained by the flies carrying different alleles at the *Ten-a* gene, altering the function of a specific subcircuit within the central complex of the brain. This illustrates how polymorphisms in a single gene can result in genetic variance heterogeneity for a complex behavioral trait.

### Linkage-disequilibrium can lead to genetic variance heterogeneity in a population

Genetic variance heterogeneity can be observed in a population where two, or more, alleles that have different effects on a trait are linked. The reason for this is as follows. A common assumption when estimating the effect of a particular locus on a trait, regardless of whether it is on the mean or the variance, is that it is bi-allelic. When analyzing a population it is, however, not possible to measure the effect closely linked functional polymorphisms independently. This as the test for association to a marker will capture the joint effect of all mutations that are in linkage disequilbrium (LD) with the alleles at the tested locus in the population. If there is LD between more than two polymorphisms that affect the tested trait, genetic variance heterogeneity might emerge. If it will does ultimately depend on the LD-pattern between the functional alleles, and on their phenotypic effects. To illustrate this, we will use our findings when dissecting a variance-heterogeneity locus in *A. Thaliana*. There we found that the genetic variance heterogeneity was due to an extended LD across multiple polymorphic sites near the gene *MOT1*, which all affected the plants ability to accumulate molybdenum from the soil^15^. Several marker alleles across this locus were in LD with three different functional alleles: two that increased and one that decreased the molybdenum concentration in the leaves when compared to the major alleles at these three loci that were in LD with the alternative alleles at the markers. The plants that carry marker-alleles, which tags the three functional alleles that either increase or decrease the phenotype, thus have a more variable phenotype than the plants with the opposite marker-alleles. The resulting variance heterogeneity was very strong, with a sevenfold difference in phenotypic variance between the groups of accessions that carried the alternative alleles.

The two examples above illustrate two empirically revealed genetic mechanisms that can lead to genetic variance heterogeneity in a population. In the remainder of this paper, we focus on a third possible explanation – epistasis – to clarify its connection to genetic variance-heterogeneity as this has not previously been discussed in detail in the literature. This will be done by first briefly recapitulating the concept of genotype-to-phenotype (GP) mapping across multiple loci, and then use this as a tool to illustrate how different types of theoretical and empirically evidenced genetic interactions (epistasis) will, or will not, lead to genetic variance-heterogeneity at the interacting loci. This will provide a deeper insight into how these two genetic concepts are related.

## Results

For a complex trait, affected by many alleles across multiple loci, the genotype to phenotype (GP) space, i.e. the phenotype produced by every possible genotype, can be extremely large. This is because the number of possible multi-locus genotypes, the full space of genotypic possibilities, grows exponentially with the number of loci that regulate the trait. In the presence of epistasis, each of these multi-locus genotypes can in theory have their own unique phenotypic effects, giving rise to an almost infinitely complex map from genotype to phenotype. In practice, it is difficult to empirically characterize more than small fraction of this genotypic space. Partially for practical reasons, genetic studies therefore often either ignore genetic interactions completely to focus on the marginal effect of contributing loci, or focus on a smaller subset of the possible multi-locus genotypes.

### Variance-heterogeneity as an emerging property of epistatic gene-action

The marginal additive effect of a locus is the change in the phenotype due to an allele-substitution at this locus, averaged across all genetic backgrounds in the population. It can be thought of as a projection from the multi-dimensional GP-space, down to one dimension. Fig. 2 illustrates this for a theoretical GP-space involving only two loci A and B. In this example, locus B capacitates (turn on) the effect of locus A, so that A displays a phenotypic effect only when combined with the allele B_2_. The result is that under many allele frequencies, locus A will display a substantial marginal effect, but locus B will not. The marginal effect displayed by locus A is “diluted” compared to its full potential effect on the phenotype of an individual carrying it. It might, however, still be large enough for the locus to contribute substantial additive genetic variance in a population, as there will be a mean difference in the trait between the groups of individuals that carry the alternative alleles. It is also worth noting that while locus B does not have a measurable marginal effect on the phenotypic mean, there is a difference in variance between the genotypes. Thus, in this particular example, locus A does not display genetic variance heterogeneity, but locus B does. As a result, locus A might be detected in a conventional Genome Wide Association (GWA) or Quantitative Trait Locus (QTL) analyses for additive marginal effects, but locus B will not. Locus B might however be detected in an analysis looking specifically for genetic variance heterogeneity^11,13^.

**Figure 2.**
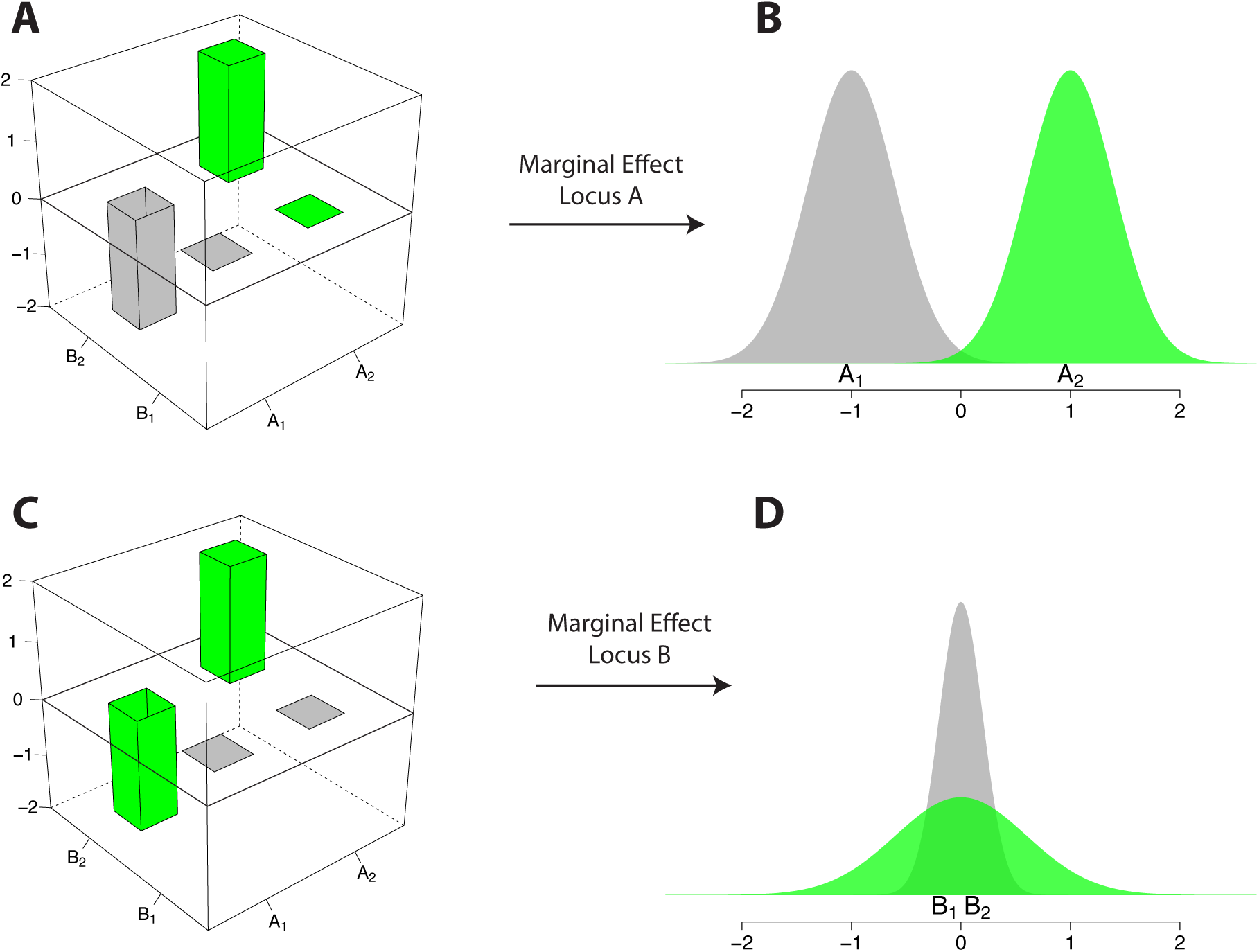
Theoretical example of pairwise capacitating epistatic gene action leading to a marginal additive effect for one, and a variance heterogeneity effect for the other, interacting locus. The alternative alleles at the two loci A and B in a haploid, or inbred, organism are denoted A_1_/A_2_ and B_1_/B_2_, respectively. Panels **A** and **C** show the phenotype associated with each of the four genotypes, i.e. the full GP-space. Panels **B** and **D** show the marginal distributions of the phenotype for the different alleles at loci A and B, obtained by comparing individuals with one allele (grey) to individuals with the opposite allele (green) at the respective loci. Locus B capacitates (turn on) the effect of locus A, so that A displays a phenotypic effect only when combined with the allele B2. Because of this, locus A displays a marginal effect on the phenotypic mean, but not on the variance, when the allele frequencies are 0.5 for the alleles at both loci. Locus B displays no marginal mean effect, but an effect on the phenotypic variance, i.e. genetic variance heterogeneity, when the allele frequencies are 0.5 for the alleles at both loci (**D**).

Fig. 3 presents another theoretical GP-space involving two loci. In this example, the genotype A_2_B_2_ has an effect of two on an arbitrarily chosen phenotypic scale, whereas the other three genotypes have an effect of zero. The result is that both loci display a marginal effect on both the mean and the variance. Just like in the previous example (Fig. 2), the marginal effect on the mean is diluted compared to its full effect on the phenotype of an individual carrying it. Under most allele frequencies, both loci will however contribute additive genetic variance. Conventional GWA and QTL analysis methods, as well as analyzes looking for marginal effects on the phenotypic variability, might in cases like this identify the two loci. Which of the two analysis approaches, looking for mean or variance effects, that has the best power will depend on the allele frequencies at the two loci.

**Figure 3.**
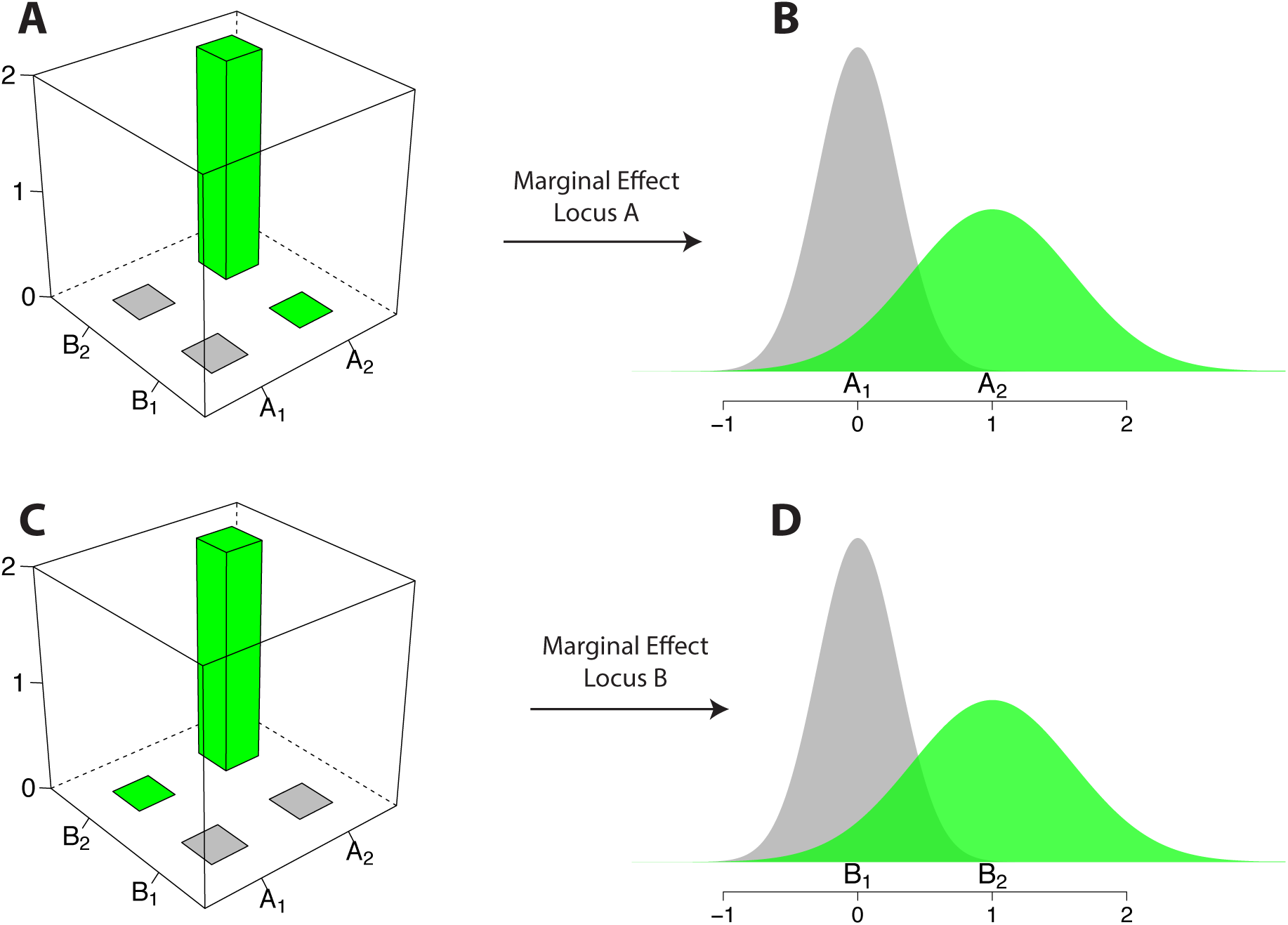
Theoretical example of pairwise epistatic gene action leading to marginal additive and genetic variance heterogeneity effects at both interacting loci. The alternative alleles at the two loci A and B in a haploid, or inbred, organism are denoted A_1_/A_2_ and B_1_/B_2_ respectively. Panels **A** and **C** show the phenotype associated with each of the four genotypes, i.e. the full GP-space. Panels **B** and **D** show the marginal phenotypic distributions of locus A and B, obtained by comparing individuals with one allele (grey) to individuals with the opposite allele (green) at the respective loci. With this underlying Genotype-Phenotype space, both loci will display both marginal additive (mean) and variance heterogeneity effects when the allele frequencies are 0.5 for the alleles at both loci.

Fig. 4 illustrates a theoretical GP-space where the direction of the phenotypic effect of an allele is completely reversed depending on the genetic background at the other locus. When combined with the B_1_ allele, A_1_ increases and A_2_ decreases the phenotype. When combined with the B_2_ allele, the effects are reversed. When estimating the marginal effect of a locus in such a GP-space the averaging across genetic backgrounds will lead to much, or all, of the phenotypic effect being canceled. At certain allele frequencies, 50% in this case, none of the loci will display any marginal effect on the mean or the variance. They will therefore not be detectable by their marginal effects, regardless of whether the scan is looking for effects on the mean or the variance of the trait. In order to identify the two loci as important contributors to the phenotype, alternative analysis methods such as a two dimensional scan for epistatic interactions^16,17^ are needed.

**Figure 4.**
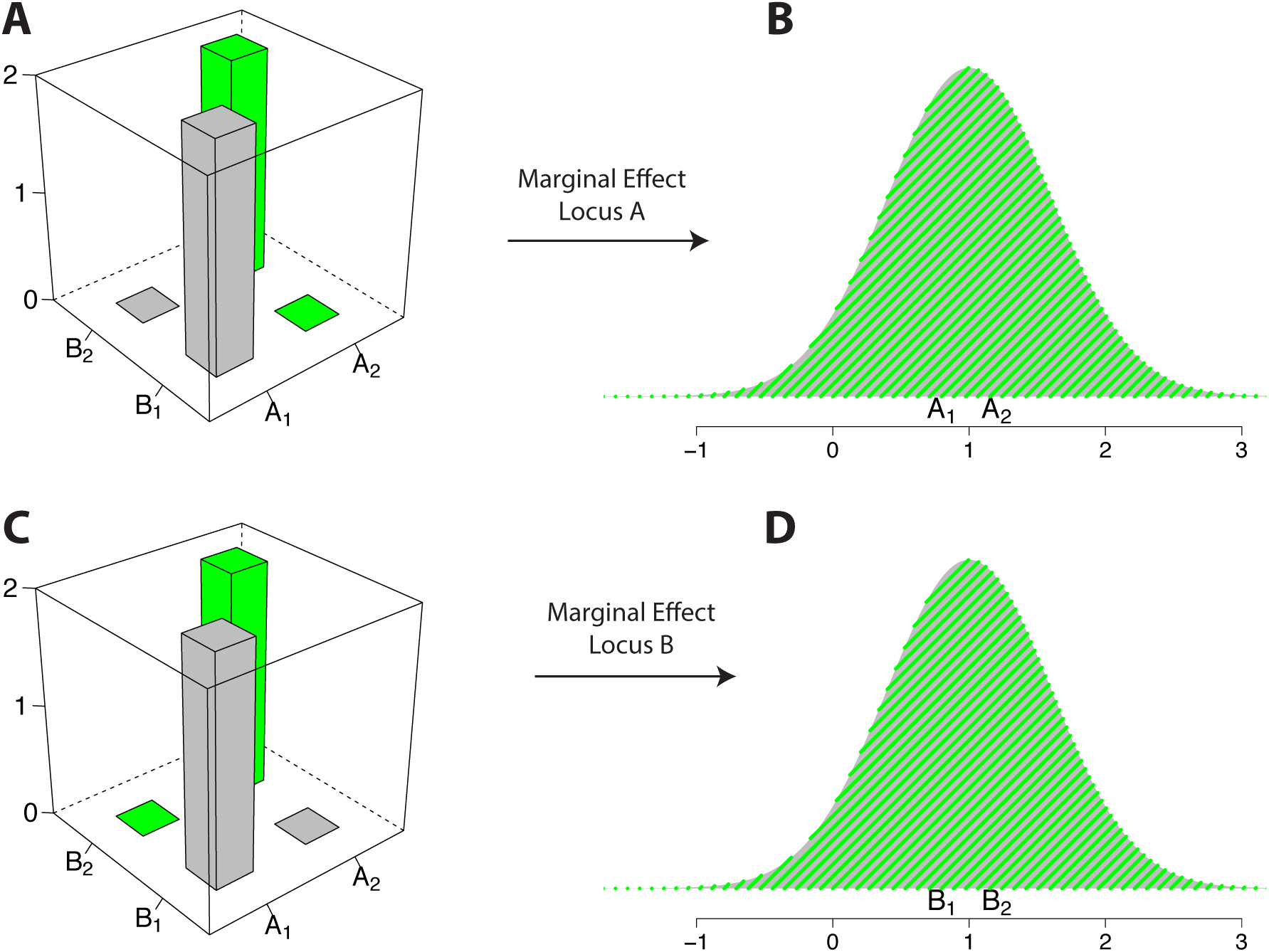
Theoretical example of pairwise epistatic gene action where no marginal additive or variance-heterogeneity effects are observed. The alternative alleles at the two loci A and B in a haploid, or inbred, organism are denoted A_1_/A_2_ and B_1_/B_2_ respectively. Panels **A** and **C** show the phenotype associated with each of the four genotypes, i.e. the full GP-space. Panels **B** and **D** show the marginal phenotypic distributions of locus A and B, obtained by comparing individuals with one allele (grey) to individuals with the opposite allele (green) at the respective loci. The direction of the phenotypic effect is completely reversed, depending on the genetic background at the other locus. Because of this, none of the loci display any marginal additive or variance heterogeneity effect when the allele frequencies are 0.5 for the alleles at both loci.

### Multi-locus epistasis and variance-heterogeneity

Unlike the theoretical examples in Fig. 2-4, most complex traits are affected by more than two loci. This means that even if one considers two-locus epistasis, this will in practice likely be a simplification of the true GP-space for the studied trait. In mathematical terms, one would be studying a projection from the high dimensional GP-space, down to a lower set of dimensions. But as with the marginal effects, this lower dimensional “shadow” might provide important insights to the genetic regulation of the trait. It can for instance facilitate the identification of causal alleles at multiple loci, as well as reveal functional dependencies between them.

In a recent study, we re-analyzed a large population of haploid yeast segregants to find a strong connection between high-order epistasis and variance-heterogeneity at the individual interacting loci^18^. In short, the size of the experimental population allowed us to fully characterize GP-spaces of up to 6 loci, consisting of 2^6^ = 64 genotypes, for multiple traits. Using these, we could evaluate how epistatic gene action contributed to the multi-locus GP-space and also identify how many of the multi-locus genotypes gave rise to phenotypes far from additive expectations. In particular, we identified several cases of capacitating epistasis where certain loci acted by moderating (i.e. turning on and off) the phenotypic effects of many other loci. Despite epistatic gene-action being common in the high-order GP-spaces, the additive genetic variance (V_A_) was much larger than the epistatic variance for all of the analyzed traits, illustrating how V_A_ can be an emergent property from epistatic gene action. These multi-locus GP-spaces can also be used to illustrate how marginal additive and variance-heterogeneity effects emerged from epistatic gene-action. An example from the analyses of this yeast population is provided in Fig. 5. There, we show the GP-space, the phenotype associated with every possible genotype, of 6 QTLs that regulate yeast growth in manganese sulfate containing growth-medium. One of the 6 QTLs capacitates the effect of the other 5, i.e. it is an empirical multi-locus example of what was theoretically illustrated in Fig. 2. Due to this capacitating effect (Fig 5A), the capacitor QTL displays pronounced genetic variance heterogeneity (Fig 5B). The other 5 QTLs display much lower levels of genetic variance heterogeneity (Fig 5C and D).

**Figure 5.**
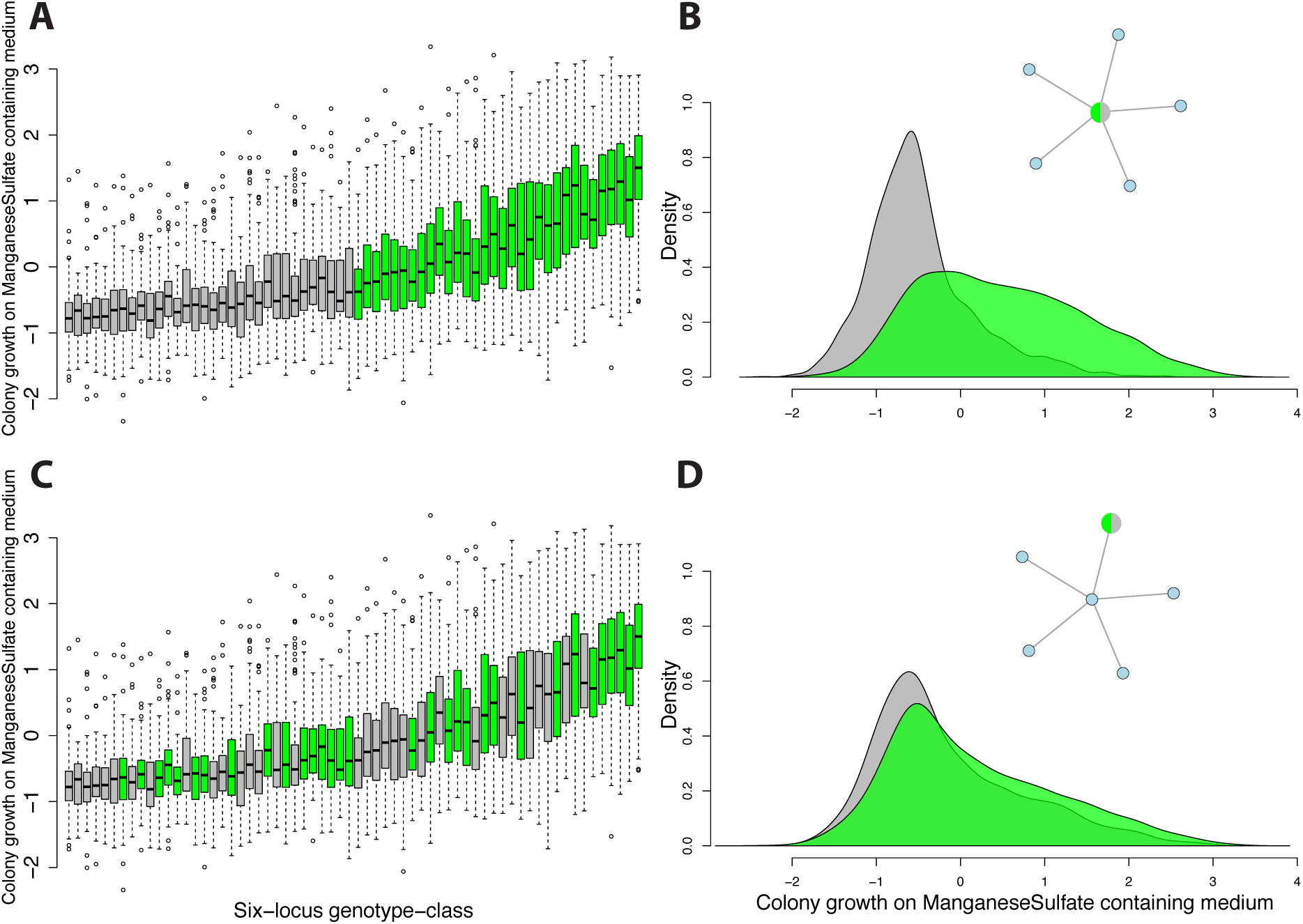
High order epistasis regulating growth in yeast leads to genetic variance-heterogeneity at the interacting loci. An epistatic network involving multiple QTLs regulates the growth of yeast colonies on media containing manganese sulfate (full details on the analysis done to identify this network is available in^18^). The full GP-space of a 6-locus network of these QTLs, made up of 2^6^ = 64 genotypes, is shown in panels **A** and **C**. The interactions in the epistatic network are illustrated in panels **B** and **D** by circles corresponding to QTLs and connections to pairwise interactions. Each boxplot in panels **A** and **C** shows the phenotype associated with one multi-locus genotype, as in Fig. 2-4, but with added information about the variability within each genotype class. One QTL in the epistatic network (large green/grey circle in the network in panel **B**) capacitates the phenotypic effects of the other QTLs (blue circles in network in **B**). Panel **B** then shows the marginal phenotypic distributions (in grey and green) for the groups of yeast segregants carrying the alternative alleles at this capacitor locus. Panel **D** shows the marginal phenotypic distributions (in green and grey) for the groups of segregants carrying alternative alleles at one of the other interacting, non-capacitor QTL (green/grey large circle in network). The capacitor locus displays a strong marginal variance heterogeneity effect (**B**), whereas the non-capacitor locus displays mostly a marginal additive effect on the phenotypic mean (**D**).

The examples above together illustrate that how epistatic gene can cause genetic variance heterogeneity and that the increased variance is the result of alleles at one locus moderating the effects of alleles at another locus. The genetic variance heterogeneity is here a “side effect” or “emergent property” of epistasis. It arises when studying the marginal effects of alleles when one should really be estimating their joint effect with alleles at other loci^18^. Screens of the genome looking for loci displaying genetic variance heterogeneity could therefore be a potential “short cut” to detect epistasis^11,19–21^, but as illustrated above not all types of epistatic gene action will lead to genetic variance heterogeneity. This strategy to detect epistatic loci can therefore not be expected to reveal all interacting loci in the genome.

## Discussion

The full GP-space underlying high-level biological traits is likely to be too complex to be directly studied in most species and populations. Genetic analyses will therefore need to rely on studies of different marginal effects of the contributing loci, emerging from the functional effects of alleles in the multi-locus genotypes of the true GP-space. When interpreting such marginal effects, for example from GWA or QTL analyses, it is important to be aware that they will often be of limited use for making direct inferences about the functional effects of individual alleles in the GP-space. Marginal effects can be very misleading in relation to the effects of alleles on the phenotypes of individuals carrying them. It is important to keep this in mind, especially if the aim of a genetic study where only marginal effects can be revealed intends to use these for prediction of individual phenotypes, such as in precision breeding or personalized medicine. When traits are regulated by genetic interactions the discrepancy between the GP-space and the marginal effects can be large. We illustrated this in Fig. 2-5, which shows that in many cases it might be impossible to make correct inferences about the contributions of a locus to the expression of a trait based on its marginal effect.

In the examples above we illustrate how some, but not all, types of epistatic gene-action can lead to marginal genetic variance-heterogeneity at the interacting loci. Further, we also show how it can emerge from other genetic mechanisms than epistasis. A genetic variance-heterogeneity signal can thus not immediately be used to identify the genetic mechanisms underlying it. But despite this, it does provide valuable information for researchers that want to interpret results from a genetic study. When a locus displays genetic variance heterogeneity, it indicates that further explorations are needed to evaluate which of the possible explanations for this signal causes it, and interpret other estimated genetic effects, such as additivity or two-way epistasis, in this context. This as it is likely that, for example, estimates of additive effects reported for loci with genetic variance heterogeneity might be sensitive to genetic background, allele-frequencies and LD-patterns among the studied individuals^15,18^.

Genome-wide screens for epistasis requires large samples, and the need for extensive multiple-testing corrections decreases power further. One-dimensional genome-scans for marginal variance-heterogeneity effects have therefore been suggested as an alternative approach to detect epistatic loci^11,13,14,21^. We have here illustrated that some types of epistatic gene-action will produce marginal variance-heterogeneity effects at the interacting loci, but also that other interacting loci might not display such a signal. Analyses of genetic variance heterogeneity can therefore not replace full epistatic analyses to reveal all interactions that contribute to a trait. They can however be an important indication that the genetic architecture of a studied trait needs to be explored beyond the marginal effects.

As illustrated in Fig. 2 and 5, genetic capacitors will often display high-levels of genetic variance heterogeneity. As has been discussed in detail elsewhere, capacitating epistasis is a mechanism contributing hidden, or cryptic, genetic variation in a population^22–25^. This as many alleles might be silently segregating in a given population, sometimes having their effects suppressed by genetic capacitors (Fig. 5), to only display their phenotypic effects upon certain changes in the genetic background. Signals of genetic variance heterogeneity might thus indicate that the population harbors hidden genetic potential that is not currently showing its phenotypic effect. Searching for such signals can therefore also be a valuable tool to identify and study cryptic genetic variation.

We have here theoretically and empirically illustrated the connection between genetic interactions (epistasis) and genetic variance heterogeneity. The two concepts are often, but not always, related and we have discussed their implications in relation to mapping and interpretation of genetic effects in genome-wide studies. We believe that by looking for genetic variance heterogeneity, valuable information can be gained in studies aiming to dissect the genetic architectures of complex traits.

## Acknowledgements

This study was funded by a grant from the Swedish Research Council FORMAS (Dnr 2013-450) to ÖC.

## Abbreviations

GP-space: Genotype to Phenotype space
QTL: Quantitative Trait Locus
GWA: Genome Wide Association
H^2^: Broad sense heritability
h^2^: Narrow sense heritability
LD: Linkage Disequilibrium

